# Population Graph GNNs for Brain Age Prediction

**DOI:** 10.1101/2020.06.26.172171

**Authors:** Kamilė Stankevičiūtė, Tiago Azevedo, Alexander Campbell, Richard Bethlehem, Pietro Liò

## Abstract

Many common neurological and neurodegenerative disorders, such as Alzheimer’s disease, dementia and multiple sclerosis, have been associated with abnormal patterns of apparent ageing of the brain. Discrepancies between the estimated brain age and the actual chronological age (brain age gaps) can be used to understand the biological pathways behind the ageing process, assess an individual’s risk for various brain disorders and identify new personalised treatment strategies. By flexibly integrating minimally preprocessed neuroimaging and non-imaging modalities into a population graph data structure, we train two types of graph neural network (GNN) architectures to predict brain age in a clinically relevant fashion as well as investigate their robustness to noisy inputs and graph sparsity. The multimodal population graph approach has the potential to learn from the entire cohort of healthy and affected subjects of both sexes at once, capturing a wide range of confounding effects and detecting variations in brain age trends between different sub-populations of subjects.

## 1. Introduction and Related Work

The link between the prevalence of neurological and neurodegenerative disorders and abnormal brain ageing patterns (Kaufmann et al., 2019) has inspired numerous studies in brain age estimation using neuroimaging data (Franke & Gaser, 2019). The resulting brain age gaps, defined as discrepancies between the estimated brain age and the true chronological age, has been associated with symptom severity of disorders such as dementia and autism (Gaser et al., 2013; Tunç et al., 2019), and could therefore be useful in monitoring and treatment of disease.

Numerous studies exist applying machine learning algorithms to the problem of brain age estimation, typically using structural magnetic resonance imaging (MRI) and genetic data (Franke & Gaser, 2019). They tend to model healthy controls separately from individuals with brain disorders, and often develop separate models for each sex (Niu et al., 2019; Kaufmann et al., 2019) without explicitly considering potential variation in ageing patterns across different subgroups of subjects. Moreover, these studies rarely include other important brain imaging modalities such as functional MRI (fMRI) time-series data, or clinical expertise of neurologists and psychiatrists, even though a combination of different modalities has been shown to improve the results (Niu et al., 2019).

In this work, we use population graphs (Parisot et al., 2017; 2018) to flexibly combine neuroimaging as well as non-imaging modalities in order to predict brain age in a clinically relevant fashion. In the population graph, the nodes contain subject-specific neuroimaging data, and edges capture pairwise subject similarities determined by non-imaging data. In addition to controlling for confounding effects (Ruigrok et al., 2014; The Lancet Psychiatry, 2016), these similarities help to exploit neighbourhood information when predicting node labels – an approach that has successfully been applied to a variety of problems in both medical and non-medical domains (Tong et al., 2017; Wang et al., 2017; Parisot et al., 2018).

We analyse the effectiveness of population graphs for brain age prediction by training two types of graph neural networks – the Graph Convolutional Network (Kipf & Welling, 2017) and the Graph Attention Network (Veličković et al., 2018). We additionally explore the robustness of our approach to node noise and edge sparsity as a way to estimate the generalisability of the models to real clinical settings. The code is available on GitHub at https://github.com/kamilest/brain-age-gnn.

## 2. Methods

### 2.1. Brain age estimation

The *brain age* is defined as the *apparent* age of the brain, as opposed to the person’s true (or chronological) age (Niu et al., 2019). Brain age cannot be measured directly and is generally unknown; however, since it is conceptually modelled after typical ageing of a healthy brain, it can be estimated by fitting a model that predicts the *chronological* age for healthy subjects, and applying the same model to the remaining population. The prediction is then assumed to correspond to brain age and any discrepancy is attributed to the brain age gap. Further details explaining the motivation behind this approach are provided in Appendix B.

In population graph context, the model is fitted to the subset of nodes representing healthy subjects in the training set. The graph neural networks are leveraged to generate predictions for the remaining nodes of the graph. The overview of this process is shown in Figure 1.

**Figure 1:**
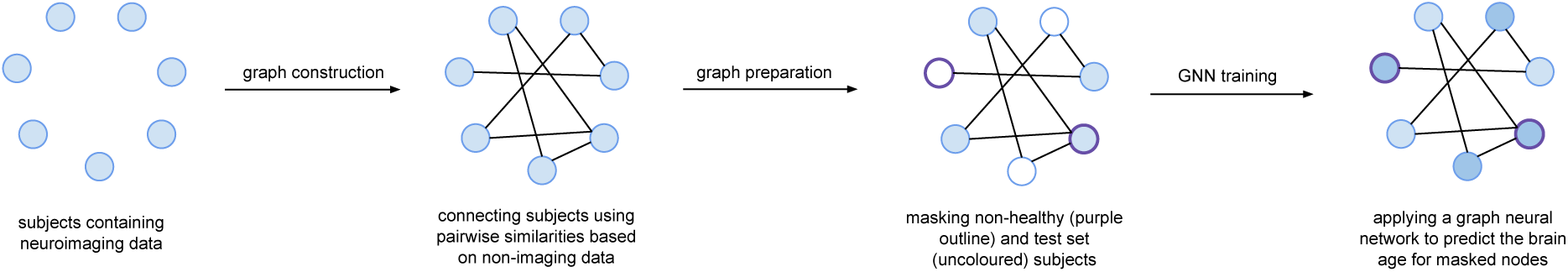
Overview of the population graph preparation and graph neural network training procedure for brain age prediction.

### 2.2. Population graphs

We combine multiple data modalities related to brain age prediction (see Section 3) using a population graph data structure, similar to the approach of Parisot et al. (2018). The nodes of the population graph correspond to individual subjects and contain features related to neuroimaging data, while the edges represent pairwise subject similarities based on non-imaging features. The edges are defined using a similarity function that computes a similarity score, and can be used to connect the patients based on the confounding effects that relate them. For simplicity, we compute the similarity scores by taking the average of indicator functions (one for each non-imaging feature), but alternative combinations (especially those which could incorporate the domain expertise of neurologists and psychiatrists) are possible. An edge is added to the graph if the similarity score is above the predefined similarity threshold. A more formal definition of population graphs is presented in Appendix A.

### 2.3. Training procedure

We train two types of graph neural network (GNN) architectures to predict brain age from population graphs, namely the Graph Convolutional Network (GCN) (Kipf & Welling, 2017), which is based on computing the graph Laplacian, and the Graph Attention Network (GAT) (Veličković et al., 2018), which operates in the Euclidean domain.

We use 10% of the dataset of healthy and affected subjects as a hold-out test set, stratifying by age and sex. The remaining data are split into five stratified cross-validation folds (with 90% training and 10% validation nodes) for model selection. In order to learn the brain age using chronological labels (as discussed in Section 2.1), we hide (or mask) the nodes of non-healthy subjects. The models are trained in a semi-supervised manner: while both the training set and the masked nodes are included in the graph, only the training node labels are visible, with the goal to learn the labels for the remaining nodes (Kipf & Welling, 2017). After the model has converged (minimising MSE loss on validation sets with early stopping), every node in the population graph (including the test set and masked nodes) has its brain age prediction.

## 3. Dataset

We use the The United Kingdom Biobank (UKB) (Sudlow et al., 2015), a continuous population-wide study of over 500,000 participants containing a wide range of measurements. We selected the UKB participants with available structural and functional magnetic resonance imaging (MRI) data, a total of 17,550 subjects.

### 3.1. Neuroimaging features

Structural MRI is used to analyse the anatomy of the brain. We use cortical thickness, surface area and grey matter volume, extracted from structural MRI images using the Human Connectome Project Freesurfer pipeline (see Glasser et al. (2013) for further discussion). We additionally experimented with the resting state functional MRI data, which approximates the activity of the brain regions over time; however, due to high computational cost this modality was excluded from training. Finally, we use the Euler index (Rosen et al., 2018) quality control metric in order to correct for any scan quality-related bias. The neuroimaging data of every subject, parcellated^1^ with Glasser parcellation (Glasser et al., 2016), were concatenated and used as node features in the population graphs.

### 3.2. Non-imaging features

Non-imaging data refers to all subject data that do not come from structural MRI and fMRI scans. We included such features as subjects’ binary (biological) sex and brain health-related diagnoses, as these variables might have a confounding effect on structural and functional connectivity of the brain (Ruigrok et al., 2014), and consequently affect the brain age (see Appendix C for further discussion of non-imaging feature selection). The non-imaging data are used to compute the inter-subject similarity scores and determine the edges of the population graph.

## 4. Results

### 4.1. Evaluation metrics

We evaluate the predictive power of the models based on their performance on the healthy subjects in the test set, for which the labels had been invisible at the training stage. We use MSE as the loss function to be optimised, and for evaluation we use Pearson’s correlation *r* and coefficient of determination *r*^2^ (Niu et al., 2019).

### 4.2. Test set performance

Considering the large size of the UKB dataset and that retraining the model using more data but without early stopping might not improve generalisation, we provide all cross-validation scores for test set evaluation instead of retraining on the entire training set and deriving a point estimate. Table 1 gives the performance metrics on the hold-out set.

**Table 1:** Test set performance of GCN and GAT models (over the five early stopping folds of the training set).

| Model | MSE | $r$ | $r^2$ |
| --- | --- | --- | --- |
| GCN1 | 28.045 ± 0.595 | 0.675 ± 0.008 | 0.445 ± 0.010 |
| GAT1 | 27.543 ± 0.758 | 0.670 ± 0.005 | 0.477 ± 0.008 |

To ensure that any variation is due to the experimental setup and not the model weights or distributions of subjects across the folds, in the following sections we consider only one fold for each graph neural network architecture. The fold was selected arbitrarily to be the first one returned by the stratified splitting procedure.

### 4.3. Robustness to population graph node feature noise

A desirable property for real-world machine learning models is their robustness to the noise and inconsistency in input data. For population graphs trained on graph neural networks, this could be estimated by adding noise to an increasing proportion of nodes.

#### Experimental setup

An increasing proportion of population graph nodes is corrupted by randomly permuting their features. Then the model is retrained and tested on the hold-out test set, measuring the change in performance. To make sure that any effect on the evaluation metrics is due to added noise and not the changing dataset splits, the model is trained on a single dataset split while the noise is added to different subjects. Moreover, to ensure that the effect on test set performance is due to the interaction with neighbourhoods and not due to the individual node features, only the nodes in the training set are corrupted. For each of the GCN and GAT models, we repeat the experiment five times.

#### Results

The impact of node feature corruption on the *r*^2^ of the GNN models is shown in Figure 2 (the behaviour for *r* is similar, see Appendix D). For both architectures the performance decreased as more training nodes were corrupted, and dropped drastically when more than half of the training nodes had their features permuted.

**Figure 2:**
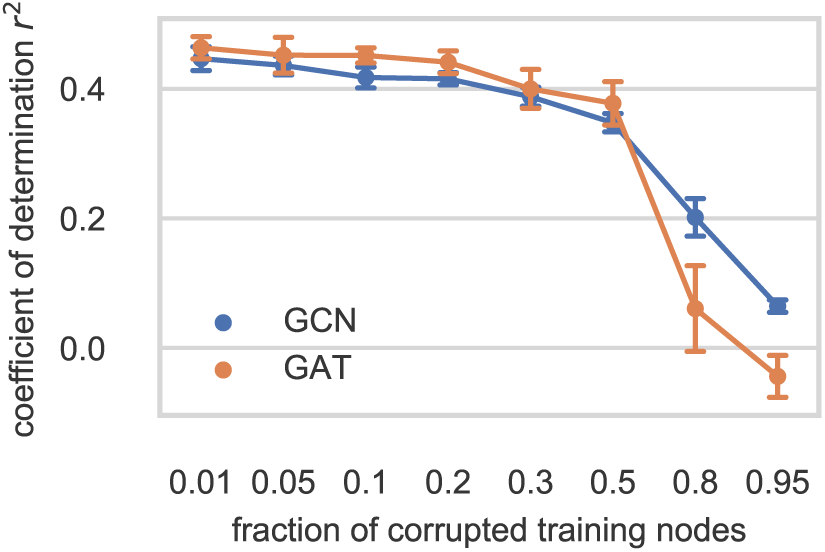
The effect of permuting node features on *r*^2^ of the test set, with error bars representing one standard deviation.

### 4.4. GNN dependence on population graph topology

#### Experimental setup

The assumption behind the population graph model is that the edge structure helps to control for confounding effects and provides additional information for brain age prediction. We test this by removing an increasing proportion of edges from the population graph, repeating the procedure five times using a different random seed. The more edges are removed, the less neighbourhood structure the graph neural network models can exploit, having to rely on individual node features.

#### Results

The effect of increasing edge sparsity on predictive power of the GNN models is shown in Figure 3. Compared to the results of the previous experiment, where *r*^2^ drastically dropped with increased noise, the loss of information contained in edges and neighbouring nodes did not affect the performance of the models. We infer this from the wide standard deviation intervals that overlap across almost all edge loss levels.

**Figure 3:**
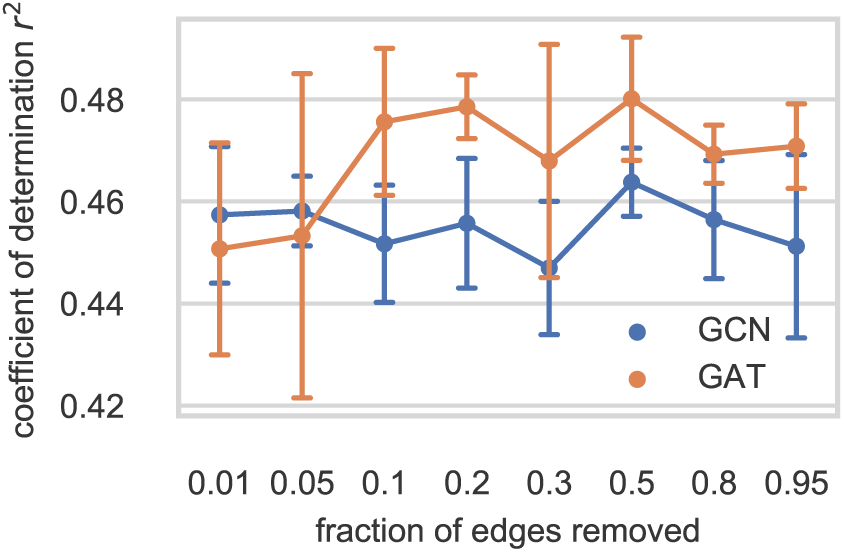
The effect of removing edges on *r*^2^ of the test set, with error bars representing one standard deviation.

## 5. Discussion

In literature on brain age estimation, many alternative models perform better than the proposed GNN models, including an XGBoost model in Kaufmann et al. (2019) with *r* = 0.93 (female) and *r* = 0.94 (male), a Gaussian process regression model in Cole et al. (2018) with *r* = 0.94, *r*^2^ = 0.88, and similar results in a variety of models using the BrainAGE technique, summarised in Franke & Gaser (2019). However, these approaches often eliminate important confounding (e.g. sex and brain health) effects by fitting separate models, use very small (i.e. a few hundred people) and consequently less diverse datasets, and filter out low-quality scans. While this improves the performance, it might affect the applicability of these models to real clinical settings, where data quality is less consistent.

The node feature noise experiment shows that high levels of node corruption in the training set could drastically worsen the predictions for the uncorrupted test nodes. This result is expected as not only does the noise propagate to neighbourhoods affecting individual predictions, but there is also less useful training data available for the GNN architectures to learn from.

The edge removal experiment shows that the models rely more on the features of individual nodes rather than the graph structure defined by the similarity metrics. One explanation could be that the brain age depends more on the feature interactions within a single brain rather than the more universal signs of ageing; however, it seems more likely that the similarity metrics used (and the simple averaging technique to combine them) were not informative enough to allow for effective sharing of feature and label information. For example, the work of Parisot et al. (2018), which used population graphs to achieve state-of-the-art results in brain disorder classification, shows that results can vary significantly based on the selection of similarity features alone, with up to 20% difference in mean accuracy scores. At the same time, regression tasks such as brain age prediction are also more difficult in nature compared to classification tasks.

## 6. Conclusion

In this work we have combined several imaging as well as non-imaging modalities into a population graph in order to predict the apparent brain age for a large and diverse dataset of subjects. The population graph representation allows to control for the confounding effects through pairwise similarities (i.e. the population graph edges) rather than fitting of separate models, and to train the entire dataset at once without extensive filtering of the data (i.e. conditions that are closer to real clinical settings).

Combination of multimodal data might not be as feasible or practical with alternative brain age prediction approaches (as it might be harder to logically separate and control the relative importance of the few non-imaging features among many more imaging features in the same vector), but might become more important with future advancements in neuroscience, growth of neuroimaging datasets, and growth in computational resources to support their processing. Moreover, training on more data (while mitigating memory constraints relating to both the GNNs and storing the entire dataset as a single object), incorporating additional (e.g. genetic (Parisot et al., 2018)) modalities, and trying alternative state-of-the-art graph neural network architectures could give a much better picture of the potential of this approach.

Consistent and unified processing of the different data modalities is also important (regardless of the downstream task or analysis method) as there is a widespread community effort to combat the reproducibility crisis in both neuroimaging (Gorgolewski & Poldrack, 2016) and machine learning^2^ fields caused by, among other factors, the lack of transparency in preprocessing methods and software errors due to bad software engineering practices (Poldrack et al., 2017). While the efforts to improve reproducibility in neuroimaging are currently targeted at consistent (yet separate) processing of functional and structural MRI with libraries like *fMRIPrep* (Esteban et al., 2019) or *sMRIPrep*^3^, this work, to the best of our knowledge, is one of the first to additionally incorporate non-imaging data modalities. While this pipeline was designed to prepare the data specifically for population graphs, it has sufficient flexibility to be extended to more preprocessing options, and adapted to work independently of the downstream analysis method.

## Acknowledgements

This research was co-funded by the NIHR Cambridge Biomedical Research Centre and a Marmaduke Sheild grant to Richard A.I. Bethlehem and Varun Warrier. The views expressed are those of the author(s) and not necessarily those of the NHS, the NIHR or the Department of Health and Social Care.

## Footnotes

1 A *parcellation* splits an image of a brain into biologically meaningful regions for downstream analysis, compressing per-voxel measurements into per-parcel summaries. A *voxel* is a discrete volumetric element.

2 https://reproducibility-challenge.github.io/neurips2019/

3 https://github.com/poldracklab/smriprep

## References

Brayne, C., Ince, P. G., Keage, H. A., McKeith, I. G., Matthews, F. E., Polvikoski, T., and Sulkava, R. Education, the brain and dementia: neuroprotection or compensation? eclipse collaborative members. Brain, 133(8): 2210–2216, 2010.

Cole, J., Ritchie, S., Bastin, M., Valdés Hernández, M., et al. Brain age predicts mortality. Molecular Psychiatry, 23 (5):1385–1392, 2018.

Dukart, J., Schroeter, M. L., Mueller, K., Initiative, A. D. N., et al. Age correction in dementia–matching to a healthy brain. PloS one, 6(7), 2011.

Esteban, O., Markiewicz, C. J., Blair, R. W., Moodie, C. A., Isik, A. I., Erramuzpe, A., Kent, J. D., Goncalves, M., DuPre, E., Snyder, M., et al. fMRIPrep: a robust preprocessing pipeline for functional MRI. Nature methods, 16 (1):111–116, 2019.

Franke, K. and Gaser, C. Ten years of BrainAGE as a neuroimaging biomarker of brain aging: What insights have we gained? Frontiers in Neurology, 10:789, 2019.

Gaser, C., Franke, K., Klöppel, S., Koutsouleris, N., Sauer, H., Initiative, A. D. N., et al. Brainage in mild cognitive impaired patients: predicting the conversion to alzheimer’s disease. PloS one, 8(6), 2013.

Glasser, M. F., Sotiropoulos, S. N., Wilson, J. A., Coalson, T. S., Fischl, B., Andersson, J. L., Xu, J., Jbabdi, S., Webster, M., Polimeni, J. R., Van Essen, D. C., and Jenkinson, M. The minimal preprocessing pipelines for the human connectome project. NeuroImage, 80:105–124, 2013. ISSN 1053-8119. doi: https://doi.org/10.1016/j.neuroimage.2013.04.127. URL http://www.sciencedirect.com/science/article/pii/S1053811913005053. Mapping the Connectome.

Glasser, M. F., Coalson, T. S., Robinson, E. C., Hacker, C. D., Harwell, J., Yacoub, E., Ugurbil, K., Andersson, J., Beckmann, C. F., Jenkinson, M., et al. A multi-modal parcellation of human cerebral cortex. Nature, 536(7615): 171, 2016.

Gorgolewski, K. J. and Poldrack, R. A. A practical guide for improving transparency and reproducibility in neuroimaging research. PLoS biology, 14(7), 2016.

Grady, C. L. and Craik, F. I. Changes in memory processing with age. Current opinion in neurobiology, 10(2):224–231, 2000.

Gray, J. R., Chabris, C. F., and Braver, T. S. Neural mechanisms of general fluid intelligence. Nature neuroscience, 6(3):316–322, 2003.

Kaufmann, T., van der Meer, D., Doan, N. T., Schwarz, E., et al. Common brain disorders are associated with heritable patterns of apparent aging of the brain. Nature Neuroscience, 22(10):1617–1623, 2019. ISSN 1546-1726. doi:10.1038/s41593-019-0471-7. URL https://doi.org/10.1038/s41593-019-0471-7.

Kipf, T. N. and Welling, M. Semi-supervised classification with graph convolutional networks. In International Conference on Learning Representations (ICLR), 2017.

Kliegel, M. and Jager, T. Delayed–execute prospective memory performance: The effects of age and working memory. Developmental neuropsychology, 30(3):819–843, 2006.

Niu, X., Zhang, F., Kounios, J., and Liang, H. Improved prediction of brain age using multimodal neuroimaging data. Human Brain Mapping, 2019.

Parisot, S., Ktena, S. I., Ferrante, E., Lee, M., et al. Spectral graph convolutions on population graphs for disease prediction. MICCAI, 2017.

Parisot, S., Ktena, S. I., Ferrante, E., Lee, M., et al. Disease prediction using graph convolutional networks: Application to Autism Spectrum Disorder and Alzheimer’s disease. Medical Image Analysis, 48:117–130, August 2018. doi: https://doi.org/10.1016/j.media.2018.06.001.

Poldrack, R. A., Baker, C. I., Durnez, J., Gorgolewski, K. J., Matthews, P. M., Munafó, M. R., Nichols, T. E., Poline, J.-B., Vul, E., and Yarkoni, T. Scanning the horizon: towards transparent and reproducible neuroimaging research. Nature reviews neuroscience, 18(2):115, 2017.

Rosen, A. F., Roalf, D. R., Ruparel, K., Blake, J., Seelaus, K., Villa, L. P., Ciric, R., Cook, P. A., Davatzikos, C., Elliott, M. A., et al. Quantitative assessment of structural image quality. Neuroimage, 169:407–418, 2018.

Ruigrok, A. N., Salimi-Khorshidi, G., Lai, M.-C., Baron-Cohen, S., Lombardo, M. V., Tait, R. J., and Suckling, J. A meta-analysis of sex differences in human brain structure. Neuroscience & Biobehavioral Reviews, 39: 34–50, 2014.

Steffener, J., Habeck, C., O’Shea, D., Razlighi, Q., Bherer, L., and Stern, Y. Differences between chronological and brain age are related to education and self-reported physical activity. Neurobiology of aging, 40:138–144, 2016.

Sudlow, C., Gallacher, J., Allen, N., Beral, V., Burton, P., Danesh, J., Downey, P., Elliott, P., Green, J., Landray, M., et al. UK Biobank: an open access resource for identifying the causes of a wide range of complex diseases of middle and old age. PLOS Medicine, 12(3):e1001779, 2015.

The Lancet Psychiatry. Sex and gender in psychiatry. Lancet Psychiatry, 3(11):999, 2016.

Tong, T., Gray, K., Gao, Q., Chen, L., Rueckert, D., Initiative, A. D. N., et al. Multi-modal classification of Alzheimer’s disease using nonlinear graph fusion. Pattern recognition, 63:171–181, 2017.

Tunç, B., Yankowitz, L. D., Parker, D., Alappatt, J. A., Pandey, J., Schultz, R. T., and Verma, R. Deviation from normative brain development is associated with symptom severity in autism spectrum disorder. Molecular Autism, 10(1):46, 2019.

Veličković, P., Cucurull, G., Casanova, A., Romero, A., Lió, P., and Bengio, Y. Graph Attention Networks. International Conference on Learning Representations, 2018. URL https://openreview.net/forum?id=rJXMpikCZ.

Wang, Z., Zhu, X., Adeli, E., Zhu, Y., Nie, F., Munsell, B., Wu, G., et al. Multi-modal classification of neurodegenerative disease by progressive graph-based transductive learning. Medical image analysis, 39:218–230, 2017.

