## Appendix for "Population Graph GNNs for Brain Age Prediction"

### A. Population graphs

A set of $N$ subjects $S$ is connected into an undirected *population graph* $G = (V, E)$, where $V$ is the set of graph nodes (with one node uniquely representing one subject, $|S| = |V|$), and $E$ is the set of edges (representing the similarity of subjects). Each node $v \in V$ is a vector containing the individual subject's neuroimaging data, whether structural, functional, or both. The edge $(v, w) \in E$ connects subjects $s_v, s_w \in S$ based on some *similarity metric* that uses the non-imaging information of the subjects to create edges between the nodes.

Defining a good similarity metric is important to account for the confounding effects on the feature vectors (e.g. the subject's sex affects the brain volume) as well as to cluster subjects into the most informative neighbourhoods. For example, here the neighbourhoods that have similar brain age gaps could be useful. If carefully defined, similarity metrics could reflect the domain expertise of neurologists and psychiatrists.

Similarity metrics are defined using a *similarity function* $\mathrm{sim}(\cdot, \cdot)$ which takes two subjects and returns the similarity score between them (the higher the score, the more similar the subjects). In this work, we use the following similarity function (more sophisticated functions are possible):

$$\mathrm{sim}(s_v, s_w) = \frac{1}{n} \sum_{i=1}^{n} \mathbf{1}[M_i(s_v) = M_i(s_w)]. \tag{1}$$

Here $\{M_1, \dots, M_n\}$ is a set of non-imaging features that are used to compute subject similarity and $\mathbf{1}[\cdot]$ is an indicator function, in this case returning a non-zero value when the values for a given non-imaging feature $M_i$ match for the two subjects $s_v$ and $s_w$. In practice, if the metric is a real number, "matching" can be defined in terms of non-imaging features being within some constant $\epsilon > 0$.

To avoid memory issues when $|E| \sim O(N^2)$ and minimise the size of the neighbourhood to only highly similar subjects, a *similarity threshold* $\mu$ is used such that

$$(v, w) \in E \iff \mathrm{sim}(s_v, s_w) \geq \mu. \tag{2}$$

### B. Brain age estimation

Formally, the brain age $y_b$ can be expressed as the sum of the known chronological age $y_c$ and the unknown *brain age gap* $\varepsilon_g$ that is defined as the discrepancy between the chronological and the brain age (Niu et al., 2019):

$$y_b = y_c + \varepsilon_g. \tag{3}$$

It is generally assumed (Franke & Gaser, 2019) that a typical healthy person has a normally ageing brain, so the brain age corresponds to chronological age:

$$y_b \approx y_c. \tag{4}$$

Our goal is to estimate brain age $y_b$ as a function $f(\cdot)$ of brain imaging feature vector $\mathbf{x}$:

$$y_b = f(\mathbf{x}) + \varepsilon_e, \tag{5}$$

where $\varepsilon_e$ is the prediction error, while the estimate of *chronological* age is instead (from Equations (3) and (5))

$$y_c = f(\mathbf{x}) + \varepsilon, \tag{6}$$

where $\varepsilon := \varepsilon_e - \varepsilon_g$ is the error term consisting of both the brain age gap and the model prediction error.

Since the brain age $y_b$ is unknown, any (semi-)supervised machine learning model can only use chronological age as a predicted variable, following Equation (6). However, if the model is trained on healthy subjects only, $f(\cdot)$ can explain both the apparent brain age *and* the chronological age with $\mathbf{x}$, since for healthy subjects $y_b \approx y_c$ (Equation (4)) and any variance in $\varepsilon$ is assumed to contain just the prediction error $\varepsilon_e$. When the same model is applied to non-healthy subjects, $f(\cdot)$ explains the chronological age assuming the brain is healthy, and any *additional* unexplained variance in $\varepsilon$ is assumed to be the brain age gap. On the other hand, if the model is trained on both healthy and non-healthy subjects at the same time, it might learn the combined confounding effects of both normal (chronological) and disease-related (brain) ageing, thus hiding the brain age gaps (Dukart et al., 2011).

An alternative method that does not restrict training data only to healthy subjects is proposed in Niu et al. (2019). However, it requires experimentally verifying (e.g. through subjects' performance in cognitive behaviour tests) that $\varepsilon$ depends primarily on the brain age gap $\varepsilon_g$ and not the brain age prediction error $\varepsilon_e$, which is out of the scope of this paper.

### C. Hyperparameter selection

This Appendix contains the hyperparameter search configuration and the hyperparameters for the best performing models, selected by the following procedure (applied separately to the GCN and GAT model families):

1. Models were ranked by ascending average MSE loss. The model with the lowest average MSE was chosen as the reference model.
2. Models whose one standard deviation interval from their MSE did not overlap with the one standard deviation interval of the reference model MSE were excluded from ranking.

Cross-validation performances of the best-scoring models selected are shown in Figure 4. The hyperparameters for each of the short-listed models are listed in supplementary material, Appendix C.

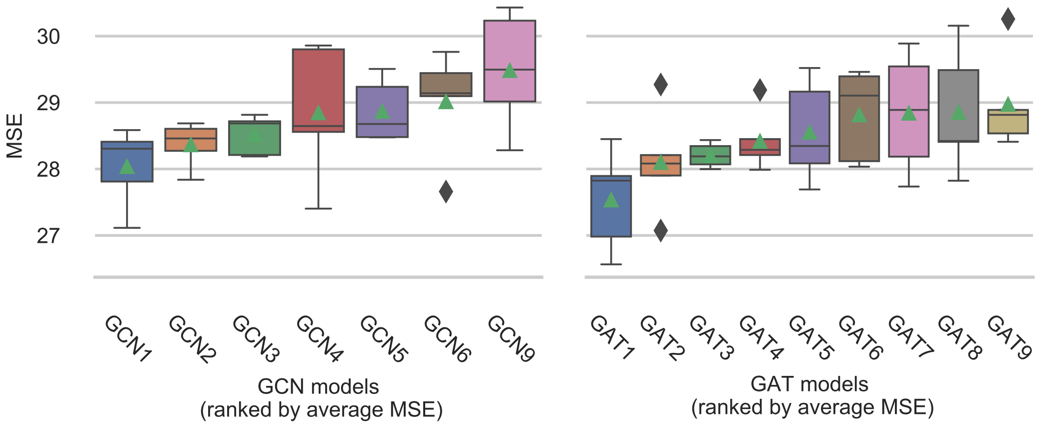


Figure 4: Highest scoring population graph and GNN parameter combinations for GCN (left) and GAT (right). The models are named according to their convolution type and ranked by ascending average MSE loss (indicated by the green triangle).

The best-ranked GCN1 and GAT1 models seemed to be the most promising and therefore have been selected for evaluation.

#### C.1. Non-imaging feature selection

Table 2 presents the non-imaging features used in this work.

#### C.2. Hyperparameter tuning configuration

The hyperparameters were searched using Bayesian optimisation strategy, and are presented in Listing 1.

[4] https://icd.who.int/browse10/2019/en

**Listing 1** Hyperparameter search configuration for the GCN and GAT model families. Similarity metrics are encoded as list of non-imaging features used along with similarity thresholds. Non-imaging encodings are presented in 2.

```
metric:
  goal: minimize
  name: cv_validation_average_mse
parameters:
  dropout:
    distribution: uniform
    max: 0.5
    min: 0
  epochs:
    value: 10000
  layer_sizes:
    distribution: categorical
    values:
    - [1024, 512, 512, 256, 256, 1]
    - [2048, 1024, 512, 256, 128, 1]
    - [1024, 512, 512, 512, 256, 256, 1]
    - [1024, 512, 512, 256, 256, 128, 128, 1]
    - [512, 512, 512, 256, 128, 1]
    - [1024, 512, 256, 128, 128, 1]
  learning_rate:
    distribution: log_uniform
    max: -2.995
    min: -9.904
  n_conv_layers:
    distribution: int_uniform
    max: 5
    min: 1
  similarity:
    distribution: categorical
    values:
    - (['SEX', 'ICD10', 'FI', 'FTE', 'MEM'], 0.8)
    - (['SEX', 'ICD10', 'FI', 'FTE', 'MEM'], 0.9)
    - (['SEX', 'FTE', 'FI', 'MEM'], 0.8)
    - (['SEX', 'ICD10', 'MEM', 'FTE'], 0.8)
    - (['SEX', 'ICD10', 'MEM', 'FI'], 0.8)
  weight_decay:
    distribution: log_uniform
    max: -2.995
    min: -9.904
```

Table 2: Summary of the non-imaging features used in this paper.

| Code | Non-imaging feature | Explanation |
|---|---|---|
| AGE | Chronological age | Used as the training label. |
| FI | Fluid intelligence score | Measures cognitive performance. Related to increased brain activity (Gray et al., 2003). |
| FTE | Years of full-time education | Associated with brain age gaps (Steffener et al., 2016) and other brain health conditions (Brayne et al., 2010). |
| ICD10 | Mental and brain health (from ICD10 diagnosis code data) | Subject mental health and nervous system disease diagnoses that might affect the structure and function of the brain (Kaufmann et al., 2019). Diagnoses were grouped by categories following the ICD10 system[4]. |
| MEM | Prospective memory result | Memory generally declines with age, and is related to changing brain activity patterns (Grady & Craik, 2000; Kliegel & Jager, 2006). |
| SEX | Binary sex (male or female) | Highly affects the size and volume of the brain (Ruigrok et al., 2014). |

#### C.3. Hyperparameters of shortlisted models

Tables 5 and 6 use encodings given in Tables 3 and 4 for similarity feature sets and layer sizes respectively.

Table 3: Similarity feature set encoding.

| Feature | FI | FTE | ICD10 | MEM | SEX |
|---|---|---|---|---|---|
| SF1 | Yes | Yes | Yes | Yes | Yes |
| SF2 | Yes | No | Yes | Yes | Yes |
| SF3 | No | Yes | Yes | Yes | Yes |
| SF4 | Yes | Yes | No | Yes | Yes |

Table 4: Layer size encoding.

| Encoding | Layer sizes |
|---|---|
| LS1 | [1024, 512, 512, 256, 256, 1] |
| LS2 | [1024, 512, 512, 512, 256, 256, 1] |
| LS3 | [1024, 512, 256, 128, 128, 1] |
| LS4 | [2048, 1024, 512, 256, 128, 1] |
| LS5 | [512, 512, 512, 256, 128, 1] |

Table 5: Shortlisted population graph and GCN model parameter combinations during the model selection process.

| Hyperparameter | GCN1 | GCN2 | GCN3 | GCN4 | GCN5 | GCN6 | GCN9 |
|---|---|---|---|---|---|---|---|
| Similarity feature set | SF1 | SF3 | SF3 | SF2 | SF2 | SF2 | SF2 |
| Similarity threshold | 0.9 | 0.8 | 0.8 | 0.8 | 0.8 | 0.8 | 0.8 |
| Layer sizes | LS1 | LS2 | LS3 | LS5 | LS3 | LS3 | LS4 |
| # convolutional layers | 5 | 3 | 1 | 2 | 5 | 3 | 4 |
| Dropout | 0.321941 | 0.042080 | 0.048596 | 0.237940 | 0.375442 | 0.386998 | 0.426491 |
| Learning rate | 0.006984 | 0.006187 | 0.005095 | 0.004731 | 0.015796 | 0.010273 | 0.003504 |
| Weight decay | 0.013118 | 0.002084 | 0.016171 | 0.002517 | 0.003114 | 0.005341 | 0.018943 |

### D. Full results

The results of node permutation and edge removal experiments are shown in Figures 5 and 6 respectively.

Table 6: Shortlisted population graph and GAT model parameter combinations during the model selection process.

| **Hyperparameter** | **GAT1** | **GAT2** | **GAT3** | **GAT4** | **GAT5** | **GAT6** | **GAT7** | **GAT8** | **GAT9** |
|---|---|---|---|---|---|---|---|---|---|
| Similarity feature set | SF2 | SF1 | SF2 | SF1 | SF1 | SF1 | SF1 | SF2 | SF2 |
| Similarity threshold | 0.8 | 0.9 | 0.8 | 0.9 | 0.9 | 0.9 | 0.9 | 0.8 | 0.8 |
| Layer sizes | LS4 | LS3 | LS3 | LS5 | LS3 | LS3 | LS3 | LS5 | LS5 |
| # convolutional layers | 2 | 2 | 3 | 2 | 3 | 2 | 3 | 3 | 3 |
| Dropout | 0.003142 | 0.306806 | 0.104624 | 0.327091 | 0.407471 | 0.323481 | 0.291117 | 0.455777 | 0.381829 |
| Learning rate | 0.013365 | 0.001679 | 0.003412 | 0.002482 | 0.003246 | 0.001462 | 0.006769 | 0.006813 | 0.003820 |
| Weight decay | 0.000605 | 0.002071 | 0.036676 | 0.001549 | 0.006715 | 0.002475 | 0.000844 | 0.001483 | 0.003226 |

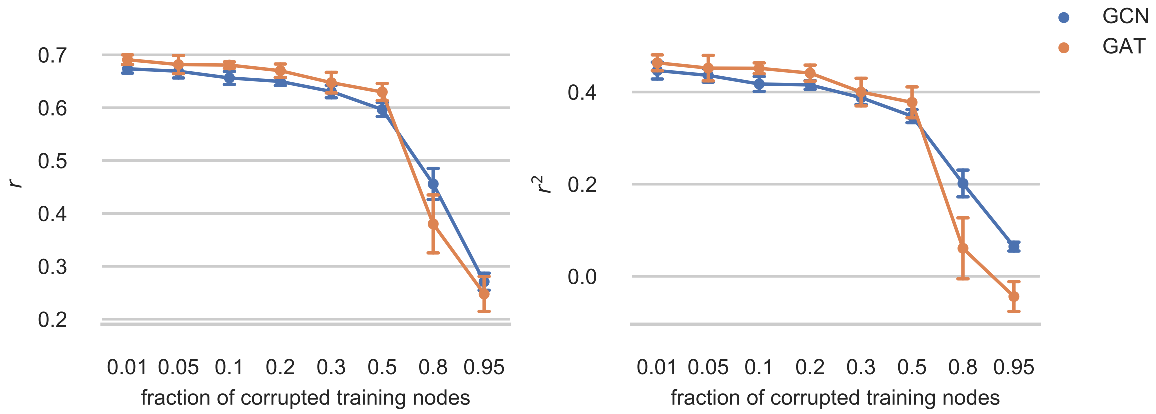


Figure 5: The effect of permuting node features on $r$ (left) and $r^2$ (right) performance metrics, with error bars representing one standard deviation.

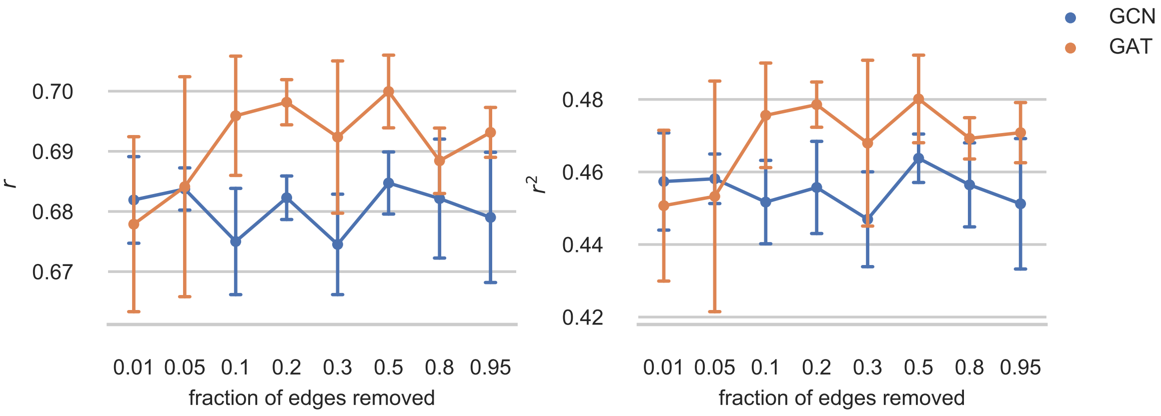


Figure 6: The effect of removing edges on $r$ (left) and $r^2$ (right) performance metrics, with error bars representing one standard deviation.
